# Field relevant variation in ambient temperature modifies the density-dependent establishment of *Plasmodium falciparum* in mosquitoes: implications for the infectious reservoir and beyond?

**DOI:** 10.1101/699850

**Authors:** Ashutosh K. Pathak, Justine C. Shiau, Matthew B. Thomas, Courtney Murdock

**Affiliations:** Dept. of Infectious Diseases, College of Veterinary Medicine, University of Georgia, Athens, GA 30602; Center for Infectious Disease Dynamics and the Department of Entomology, The Pennsylvania State University, University Park, PA16802; Odum School of Ecology, University of Georgia, Athens GA 30602; Center for Ecology of Infectious Diseases, University of Georgia, Athens GA 30602; Center for Tropical Emerging Global Diseases, University of Georgia, Athens GA 30602; Center for Vaccines and Immunology, University of Georgia, Athens GA 30602; Riverbasin Center, University of Georgia, Athens GA 30602

**Keywords:** malaria, transmission, *Plasmodium falciparum*, gametocyte density, mosquitoes, temperature, diurnal temperature, vector competence

## Abstract

The relationship between *Plasmodium falciparum* gametocyte density and infections in mosquitoes is central to understanding the rates of transmission with important implications for control. Here, we determined whether field relevant variation in environmental temperature could also modulate this relationship. *Anopheles stephensi* were challenged with three densities of *P. falciparum* gametocytes spanning a ∼10-fold gradient, and housed under diurnal/daily temperature range (“DTR”) of 9°C around means of 20°C, 24°C and 28°C. At the peak stages of infection for each temperature, the proportion of mosquitoes infected with oocysts in the midguts or infectious with sporozoites in the salivary glands were measured (referred to collectively as vector competence hereon), in addition to oocyst intensities from infected midguts. While vector competence was similar at 20 DTR 9°C and 24 DTR 9°C, the proportion of mosquitoes infected and subsequently infectious were also comparable, with evidence, surprisingly, for higher vector competence in mosquitoes challenged with intermediate gametocyte densities. For the same gametocyte densities however, severe reduction in the proportion of infectious mosquitoes was accompanied by a significant decline in vector competence at 28 DTR 9°C, although density per se showed a positive and linear effect at this temperature. Unlike vector competence, oocyst intensities decreased with increasing temperatures with a predominantly positive and linear association with gametocyte density, especially at 28 DTR 9°C. Oocyst intensities across individual infected midguts suggested temperature-specific differences in mosquito susceptibility/resistance: at 20 DTR 9°C and 24 DTR 9°C, dispersion (aggregation) increased in a density-dependent manner but not at 28 DTR 9°C where the distributions were consistently random. Limitations notwithstanding, our results have manifold implications in, for instance, how variation in temperature could modify seasonal dynamics of infectious reservoirs and transmission and the contribution of high-/patent- and low-density/sub-patent carriers, to suggestions for design and deployment of transmission-blocking vaccines/drugs, but with a cautionary note suggesting how low efficacy could lead to transmission enhancement in certain environments.

## Introduction

Targeting transmission of *Plasmodium falciparum* gametocytes to their mosquito vectors is now more pertinent than ever in light of the recent resurgence in disease incidence in sub-Saharan Africa but also for future efforts towards eliminating malaria (World Health Organization, 2018). An improved understanding of the relationship between gametocyte density in the human host and rates of transmission to the vectors is central to elucidating as well as targeting transmission and ranges from identifying individuals/sub-groups with dis-proportionately higher contributions to transmission (“infectious reservoir”) for targeted interventions, to identifying mechanisms governing transmission for new target discovery and eventual candidate selection (Rabinovich et al., 2017).

Assays of the relationship between gametocyte density and infectivity to the vector generally comprise one or more laboratory-reared vector species fed directly on an infected host or via artificial membrane, with/without serum replacement, prior knowledge of gametocyte density or genotype (Churcher et al., 2013;Stone et al., 2015;Bousema and Drakeley, 2017;Goncalves et al., 2017;Bradley et al., 2018;Grignard et al., 2018;Slater et al., 2019). Taken together, these studies suggest a saturating positive relationship between the density of gametocytes in a human host and the proportion of mosquitoes that become successfully infected, with recent evidence confirming male gametocytes as the rate-limiting factor, in line with early observations from rodent models of malaria. Thus, it is generally assumed that hosts with higher gametocyte density will infect more mosquitoes on average, which in turn shapes recommendations for interventions – reducing the density of transmission stages in the host will reduce transmission to the mosquito vector, especially since most current (e.g. artemisinin-primaquine combination therapy) and future transmission blocking interventions (e.g. vaccines) so far suggest transmission-reducing rather than transmission-blocking activity (Miura et al., 2013;Kapulu et al., 2015;Bradley et al., 2018;Colmenarejo et al., 2018;Dicko et al., 2018;Doumbo et al., 2018;Graves et al., 2018;Sagara et al., 2018). However, significant variability abounds in this relationship with the source of variability generally attributed to variation in host immuno-physiology, different mosquito species / populations, and differences across parasite genotypes in gametocyte investment. Yet, how gametocyte density within the human host influences vector processes and onwards transmission is still poorly understood - albeit with two important ramifications. First, despite the overall positive nature of the relationship between gametocyte density and mosquito infections, there is substantial uncertainty with low density (also referred to as asymptomatic / sub-patent / sub-microscopic) carriers “a third as infectious” as high-density (symptomatic / patent / microscopic) carriers (Slater et al., 2019). While identifying the causes and consequences of this uncertainty is important for determining the composition and dynamics of the infectious reservoir in the immediate future, low-density carriers are likely going to be the predominant population as we approach elimination. Second, the saturating nature of the relationship suggests that high-density carriers offer no additional benefits to transmission, which is difficult to reconcile with the evidence that rates of gametocytogenesis are under such strong selection pressure in sub-Saharan Africa (Duffy et al., 2018;Rono et al., 2018;Usui et al., 2019). However, within minutes of ingestion by the ectothermic mosquito via the host blood meal, the relative drop in temperature not only serves as the sole cue for gametogenesis (Lahondère and Lazzari, 2012; Sinden, 2015), with the parasite now subject to an environment that will substantially vary in temperature throughout the day, across seasons, and with geographic region with temperature already modulating diverse aspects of mosquito physiology, ecology, life history, and fitness in situ (Paaijmans et al., 2010;Murdock et al., 2012;Mordecai et al., 2013;Murdock et al., 2014;Johnson et al., 2015;Ryan et al., 2015;Murdock et al., 2016;Beck-Johnson et al., 2017). At its extreme, temperature can completely abrogate transmission regardless of gametocyte density (Noden et al., 1995;Eling et al., 2001;Murdock et al., 2016).

Yet, all of the studies investigating the relationship between gametocytemia and infectivity to mosquitoes to date have been performed in mosquitoes that are housed under standard laboratory conditions typically set to a single constant temperature (26°C) (Bousema et al., 2012;Churcher et al., 2013;Stone et al., 2015;Bradley et al., 2018;Stone et al., 2018;Tadesse et al., 2018;Slater et al., 2019). Variation in temperature has been shown to have strong, non-linear effects on malaria transmission, with temperatures permissive for transmission ranging from 20-30°C with an optimum around 25°C, and is the most reliable predictor of seasonal and geographical transmission rates (Mordecai et al., 2013;Johnson et al., 2015;Reiner et al., 2015;Ryan et al., 2015;Murdock et al., 2016;Shah et al., 2019). Based on this overwhelming evidence to support the influence of temperature on the parasite and vector, the current study posited that variation in field relevant factors such as temperature will shape the relationship between gametocyte density and mosquito infection, with potentially important implications for understanding the selection pressures on circulating parasite genotypes, identifying human infectious reservoirs, and evaluating the efficacy of transmission blocking / reducing interventions in variable field environments. Indeed, proof-of-concept evidence is shown in the Indian malaria vector (*Anopheles stephensi*) – *P. falciparum* system that temperature alters the relationship between density of transmission stages and infection outcomes in the mosquito vector. Female *An. stephensi* were challenged with three densities of *P. falciparum* NF54 spanning an order of magnitude. After exposure, mosquitoes were housed across three mean temperatures (20°C, 24°C and 28°C) that each had a realistic diurnal temperature fluctuation of a total of 9°C (+ 5°C / – 4°C) that span a relevant range of temperatures across which malaria is transmitted in the field (Paaijmans et al., 2010;Blanford et al., 2013;Murdock et al., 2016). The effect of gametocyte density and temperature was examined with respect to the proportion of mosquitoes that become infected and infectious, as well as overall parasite burden.

## Materials and methods

### Study design

The overall study design is depicted in Figure 1 with the indicated temperature regimes adapted from a previous study (Murdock et al., 2016). Female *An. stephensi* from the same cohort were sorted into nine groups/cups with each temperature regime receiving three cups each, 24 hours prior to the day of infection. On the day of infection, mature gametocytes of *P. falciparum* NF54 generated in vitro (Pathak et al., 2018) were serially diluted 3-fold with naïve RBCs to obtain blood-meals representing three final parasite densities with a 9-fold difference between the highest and lowest densities (∼an order of magnitude). Aliquots of the three blood-meals with the varying gametocyte density were offered to the three cups, respectively, at each temperature regime. The experiments were replicated three times with independent mosquito cohorts and parasite cultures. Since the primary objective was to determine the effect of temperature on gametocyte density and vector competence, this experimental design ensured that within each replicate, all three temperature regimes would be assessed simultaneously with the same starting parasite and vector populations in order to reduce any unexpected variations over and above those attributable to temperature. Lastly, all mosquito infections were performed between 1800 and 1900 hours which represents “dusk” on a circadian scale when Anopheline mosquitoes are most active (Rund et al., 2016).

**Figure 1:**
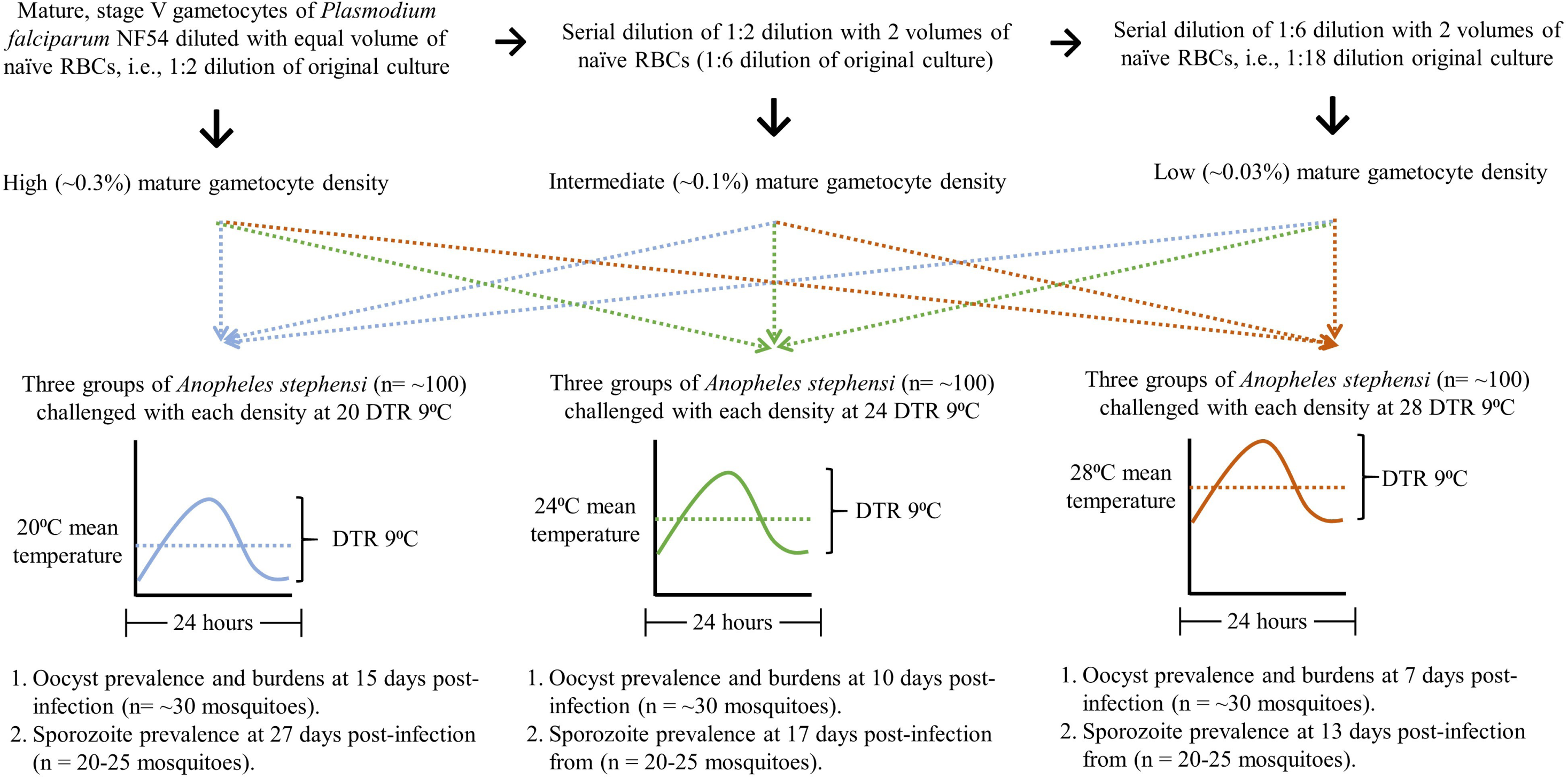
The study design.

All chemicals and consumables were purchased from Fisher Scientific Inc. unless stated otherwise.

### In vitro parasite cultures

Routine asexual cultures of *P. falciparum* NF54 and induction of gametocytogenesis in vitro were performed with cryo-preserved red blood cells (RBCs) as described in detail previously (Pathak et al., 2018). Gametocytogenesis of *P. falciparum* NF54 in vitro was monitored with Giemsa staining until 12 days post-culture, after which we assessed cultures for infectiousness through daily assays of male gametogenesis (Pathak et al., 2018). Giemsa stained slides were visualized at a 1000x magnification with an oil immersion lens while gametogenesis was assessed in vitro at 400x magnification under differential interference contrast setting on a Leica DM2500 upright microscope. Assays were performed on a 100 µl aliquot of culture re-suspended in fresh media. Infectiousness was determined in duplicate by quantifying male gametogenesis in vitro (ex-flagellation) with 10 µl culture volumes on a hemocytometer following incubation in a humidified chamber set to 24°C for 20 min. The remaining culture (∼80 µl) was concentrated by centrifugation at 1800g for two min at room temperature and Giemsa stained smears prepared as described above.

### Mosquito husbandry and experimental infections

*An. stephensi* colonies (Walter Reed Army Institute of Research, wild-type Indian strain) were housed in a walk-in environmental chamber (Percival Scientific, Perry, IA) at 27°C ± 0.5°C, 80% ± 5% relative humidity, and under a 12 hr light: 12 hr dark photo-period schedule. Adult mosquitoes were maintained on 5% dextrose (w/v) and 0.05% para-aminobenzoic acid (w/v) and provided whole human blood in glass-jacketed feeders (Chemglass Life Sciences, Vineland, NJ) through parafilm membrane maintained at 37°C to support egg production. Husbandry procedures followed methods outlined previously (Pathak et al., 2018). Briefly, eggs were rinsed twice with 1% house-hold bleach (v/v, final concentration of 0.06% sodium hypochlorite) before surface-sterilization for 1 minute in the same solution at room temperature. Bleached eggs were washed with 4-5 changes of deionized water and transferred to clear plastic trays (34.6cm L × 21.0cm W × 12.4cm H) containing 500 ml of deionized water and 2 medium pellets of Hikari Cichlid Gold fish food (HikariUSA, Hayward, CA) and allowed to hatch for 48 hours. Hatched L1 larvae were dispensed into clear plastic trays (34.6cm L × 21.0cm W × 12.4cm H) at a density of 300 larvae/1000 ml water and provided the same diet until pupation. The feeding regime consisted of 2 medium pellets provided on the day of dispensing (day 0) followed by the provision of a further 2, 4, 4 and 4 medium pellets on days 4, 7, 8 and 9 respectively. This regime allows >85% larval survival and >90% pupation within 11 days with a sex ratio of 1:1 adult males and females (unpublished observations).

Mosquito infections were performed with ∼100, 3 to 7-day old female, host-seeking *An. stephensi* sorted into nine 16/32 oz. soup cups. Three cups each were transferred to the respective temperature regimes and acclimated for ∼24 hours. On the day of infection, ex-flagellation was quantified from the cultures, in addition to gametocyte density from 3000-5000 RBCs stained with Giemsa, as described above. Parasite infected RBCs were collected into a pre-weighed 15 ml conical centrifuge tube and concentrated by centrifugation at 1800xg for 2 min at low brake setting. The media supernatant was aspirated, and weight of packed, infected RBCs estimated after subtracting the weight of the empty tube. The infected RBC pellet was then resuspended in 3 volumes of a 33% hematocrit suspension of naïve, freshly washed RBCs in human serum to achieve a hematocrit of ∼45-50%. This suspension was then serially diluted 3-fold a further two times by adding 2 volumes of naïve RBCs resuspended in human serum at 45-50% hematocrit to 1 volume of the preceding dilution resulting in three final concentrations of stage V gametocytemia (∼0.3%, 0.1%, and 0.03%) used throughout this study. This dilution scheme was developed with the objective of achieving a mature gametocytemia of ∼0.2-0.3% at the first dilution based on the gametocytemia recorded in the flasks on the day of infection and was representative of densities collated from independent studies (Adjalley et al., 2011;Miura et al., 2013;Stone et al., 2013;Miura et al., 2016;Eldering et al., 2017).

We then added an equal volume of each concentration of gametocytes to water-jacketed glass feeders maintained at ∼37°C. All nine cups of mosquitoes were placed under the respective feeders and allowed to feed for 20 min. For estimating gametocyte density, smears were prepared from the blood-meal corresponding to the 1:2 dilution for Giemsa staining. Mosquitoes were returned to the respective temperature regimes and starved for a further 48 hours to eliminate any partial or non-blood fed individuals after which they were provided cotton pads soaked in 5% dextrose (w/v) and 0.05% para-aminobenzoic acid (w/v) for the remainder of the study, as described previously (Pathak et al., 2018).

### Vector competence measurements

Vector competence was estimated at time points corresponding to peak infection intensities specific to each temperature regime with susceptibility measured as oocyst prevalence and burdens in the midguts and infectiousness as sporozoite prevalence in the salivary glands (manuscript in preparation). Specifically, midguts and salivary glands, respectively, were dissected to assess vector competence on the following days: 20 DTR 9°C, 15-17 dpi and 29 dpi; 24 DTR 9°C, 9-11 dpi and 17 dpi; and 28 DTR 9°C, 7-9 dpi and 13 dpi. At each time point, ∼25-30 mosquitoes were vacuum aspirated directly into 70% ethanol and vector competence measured as described previously (Pathak et al., 2018). Briefly, midguts were dissected, and oocysts enumerated at 400× magnification with a Leica DM2500 under DIC optics. For sporozoite prevalence, salivary glands were dissected into 5 µl of PBS, ruptured by overlaying a 22 mm^2^ coverslip and checking for presence/absence of sporozoites at either 100× or 400× magnification with the same microscope. Additionally, gravid status was also noted for each mosquito to confirm blood-feeding status.

### Data analyses

All data analyses were performed in RStudio (Version 1.1.463), an integrated development environment for the open-source R package (Version 3.5.2) (RStudio Team, 2016;R Core Team, 2018). Graphical analyses were performed with the “ggplot2” package (Wickham, 2016). Vector competence was statistically modeled using Generalized linear mixed-effects models (GLMMs) with the choice of family/distribution based on the dependent variable – 1) Oocyst and sporozoite prevalence were modeled as the probability of being infected and infectious, respectively, using a beta-binomial distribution (family=beta-binomial, link= “logit”), and 2) oocyst intensity (burdens in infected mosquito midguts only) with a negative binomial distribution (family=nbinom2, link=“log”) with the “glmmTMB” package (Brooks et al., 2017). Predictors/fixed effects comprised temperature, gametocyte density and in the case of prevalence, site of infection, i.e., midguts or salivary glands, with the relationships modeled up to three-way interactions. Temperature and site of infection were classified as categorical fixed effects while density was specified as a continuous variable exerting a linear (*x*) and quadratic (*x*^*2*^) effect on the dependent variables (Crawley, 2013). Since technical constraints meant reliable parasite counts were only available for the highest parasite concentrations (1:2 dilution), dilution was used as a proxy for gametocyte density based on the fact that all three biological replicates were performed with the same series of dilutions of the original parasite culture (1:2, 1:6 and 1:18). For all models, the random effect structure allowed for variation in the intercepts between biological replicates and/or between temperatures nested within each replicate.

The choice of family for modeling oocyst intensity was based on likelihood-based information criteria as recommended in the “bbmle” package (Bolker B and R Development Core Team, 2017), dispersion characteristics of residuals using the “DHARMa” package (Hartig, 2019), and where possible, the co-efficient of determination (“Pseudo-R-squared”) using the “sjstats” package (Lüdecke, 2018). Tests for overdispersion were performed using a predetermined threshold ratio of squared Pearson residuals over the residual degrees of freedom (overdispersion ratio= <1.5) and a Chi-squared distribution of the squared Pearson residuals with *p*-value >0.05, as described previously (Pathak et al., 2018). Once overdispersion was accounted for, the marginal means estimated by each model were then used to perform pairwise comparisons between parasite densities nested within each temperature regime, using Tukey’s contrast methods in the “emmeans” package and adjusting for multiple comparisons (Lenth, 2019).

## Results

### The effect of temperature and gametocyte density on oocyst and sporozoite prevalence

Overall, we observed a main effect of temperature on both oocyst and sporozoite prevalence: in general, oocyst and sporozoite prevalence decreased with increasing temperature, with the most notable decline at 28 DTR 9°C (Figure 2, Table 1). A non-linear quadratic (“hump-shaped”) relationship was also noted between gametocyte density and prevalence., likely driven by the patterns in oocyst prevalence at the two cooler temperatures with intermediate gametocyte density maximizing the proportion of mosquitoes infected (Figure 3, Table 1 and Supplementary table 1). While overall sporozoite prevalence largely mirrored the patterns in oocyst prevalence at 20 and 24 DTR 9°C, the overall efficiency of sporozoite establishment in the salivary glands was decreased at the warmest temperature (28 DTR 9°C; Table 1). At the highest temperature (28 DTR 9°C), we observed positive, linear effects of gametocyte density on the proportion of mosquitoes infected and subsequently infectious (Figure 3, Table 1 and Supplementary table 1). Overall, the model was able to explain 44.6% of the variation in the data with the predictors accounting for 42.7% of this fit. Pairwise comparisons of estimated marginal means from the models suggested clear differences between gametocytemia and oocyst prevalence at 28 DTR 9°C whereas at the cooler temperatures, the proportion of mosquitoes infected with oocysts was marginally higher at the intermediate relative to the lowest gametocyte densities, although, in general, none of the temperatures showed any clear differences between the two highest gametocytemias (Supplementary table 2). There was no difference in gametocytemia-dependent salivary gland prevalence at any temperature although comparisons at this site of infection should be taken with caution as data on salivary gland prevalence from one of the replicates at 20 DTR 9°C was not available for any gametocyte density (Table 1). Hence, data from the remaining two replicates was used for comparisons which may in turn impose constraints on estimating confidence intervals.

**Table 1:** Statistical models for prevalence of oocysts and sporozoites in midguts and salivary glands respectively. ^#^ For one biological replicate, sporozoite prevalence data for all three densities at 20 DTR 9°C and lowest density at 24 DTR 9°C was not available.

| **Table 1** |  |  |  |  |
| --- | --- | --- | --- | --- |
| **Predictors** (reference group= 20 DTR 9°C) | *Log-Odds* | *std. Error* | *Z value* | *p* |
| (Intercept) | 1.065 | 0.281 | 3.785 | **<0.001** |
| 24 DTR 9°C | -0.359 | 0.309 | -1.162 | 0.245 |
| 28 DTR 9°C | -1.505 | 0.328 | -4.590 | **<0.001** |
| Sporozoite prevalence (vs oocyst prevalence) | 0.148 | 0.332 | 0.447 | 0.655 |
| Gametocyte density (linear trend) | 2.116 | 1.383 | 1.530 | 0.126 |
| Gametocyte density (quadratic / “hump-shaped” trend) | -3.720 | 1.450 | -2.566 | **0.010** |
| 24 DTR 9°C * sporozoite prevalence (vs oocyst prevalence) | -0.160 | 0.430 | -0.372 | 0.710 |
| 28 DTR 9°C * sporozoite prevalence (vs oocyst prevalence) | -2.446 | 0.561 | -4.362 | **<0.001** |
| 24 DTR 9°C * Gametocyte density (linear trend) | -1.022 | 1.854 | -0.551 | 0.582 |
| 28 DTR 9°C * Gametocyte density (linear trend) | 4.099 | 2.056 | 1.993 | **0.046** |
| 24 DTR 9°C * Gametocyte density (quadratic / “hump-shaped” trend) | 0.256 | 1.963 | 0.130 | 0.896 |
| 28 DTR 9°C * Gametocyte density (quadratic / “hump-shaped” trend) | -0.047 | 2.114 | -0.022 | 0.982 |
| Sporozoite prevalence * Gametocyte density (linear trend) | 0.654 | 2.270 | 0.288 | 0.773 |
| Sporozoite prevalence * Gametocyte density (quadratic / “hump-shaped” trend) | 3.396 | 2.173 | 1.563 | 0.118 |
| 24 DTR 9°C * sporozoite prevalence * Gametocyte density (linear trend) | 0.054 | 2.983 | 0.018 | 0.986 |
| 28 DTR 9°C * sporozoite prevalence * Gametocyte density (linear trend) | -0.723 | 3.878 | -0.186 | 0.852 |
| 24 DTR 9°C * sporozoite prevalence * Gametocyte density (quadratic / “hump-shaped” trend) | 0.451 | 2.921 | 0.154 | 0.877 |
| 28 DTR 9°C * sporozoite prevalence * Gametocyte density (quadratic / “hump-shaped” trend) | -3.374 | 3.954 | -0.853 | 0.394 |
| **Random effects** |  |  |  |  |
| Random variation in intercepts between temperatures nested within each biological replicate | 0.03 |  |  |  |
| Random variation in intercepts between the three biological replicates | 0.08 |  |  |  |
| Number of observations (i.e, mosquitoes sampled) | 1469<sup>#</sup> |  |  |  |
| Marginal R<sup>2</sup> / Conditional R<sup>2</sup> | 0.427 / 0.446 |  |  |  |

**Figure 2:**
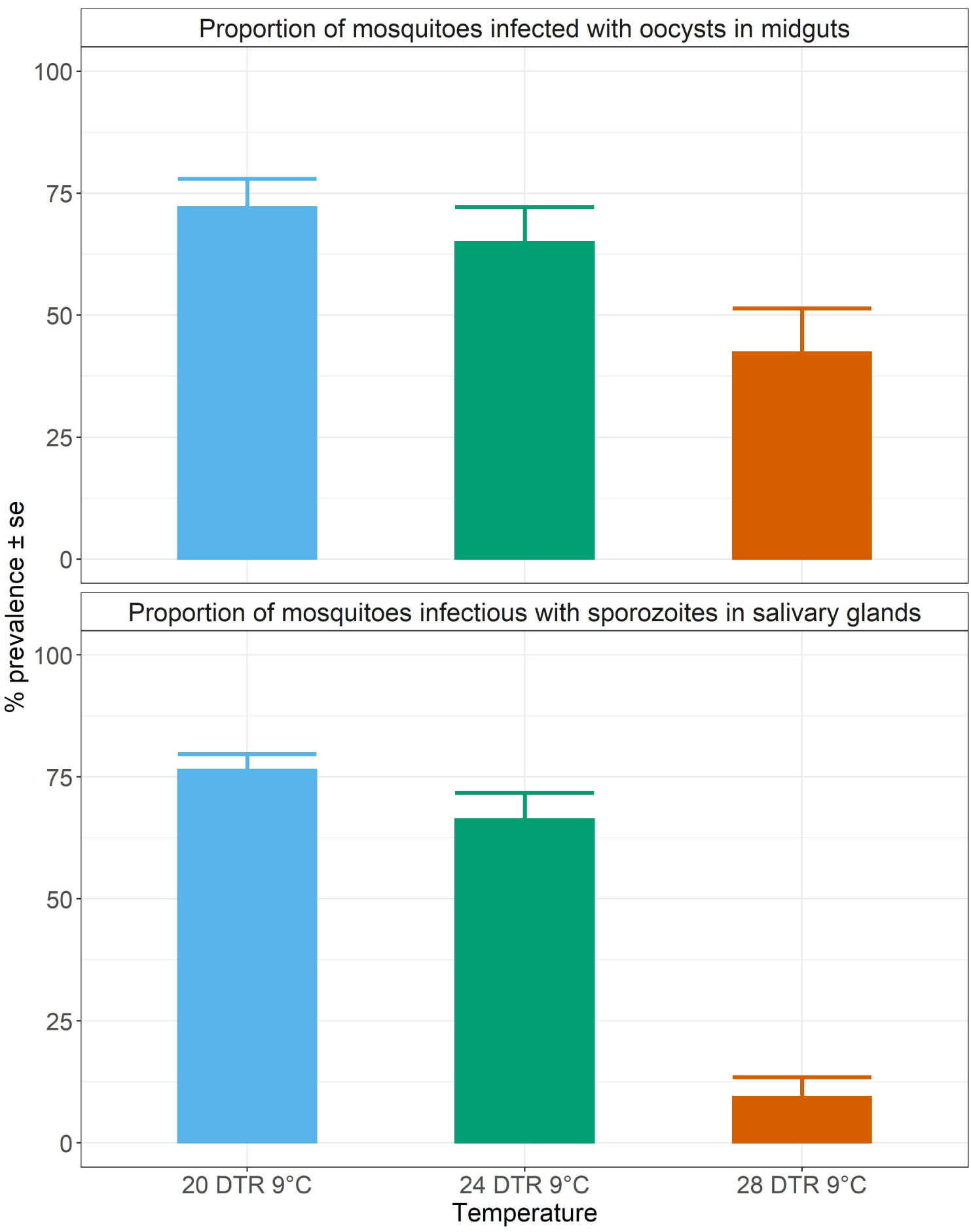
Proportion of mosquitoes infected (top pane) and infectious (bottom pane) at the three temperatures. Values represents means ± standard error (se) from three biological replicates.

**Figure 3:**
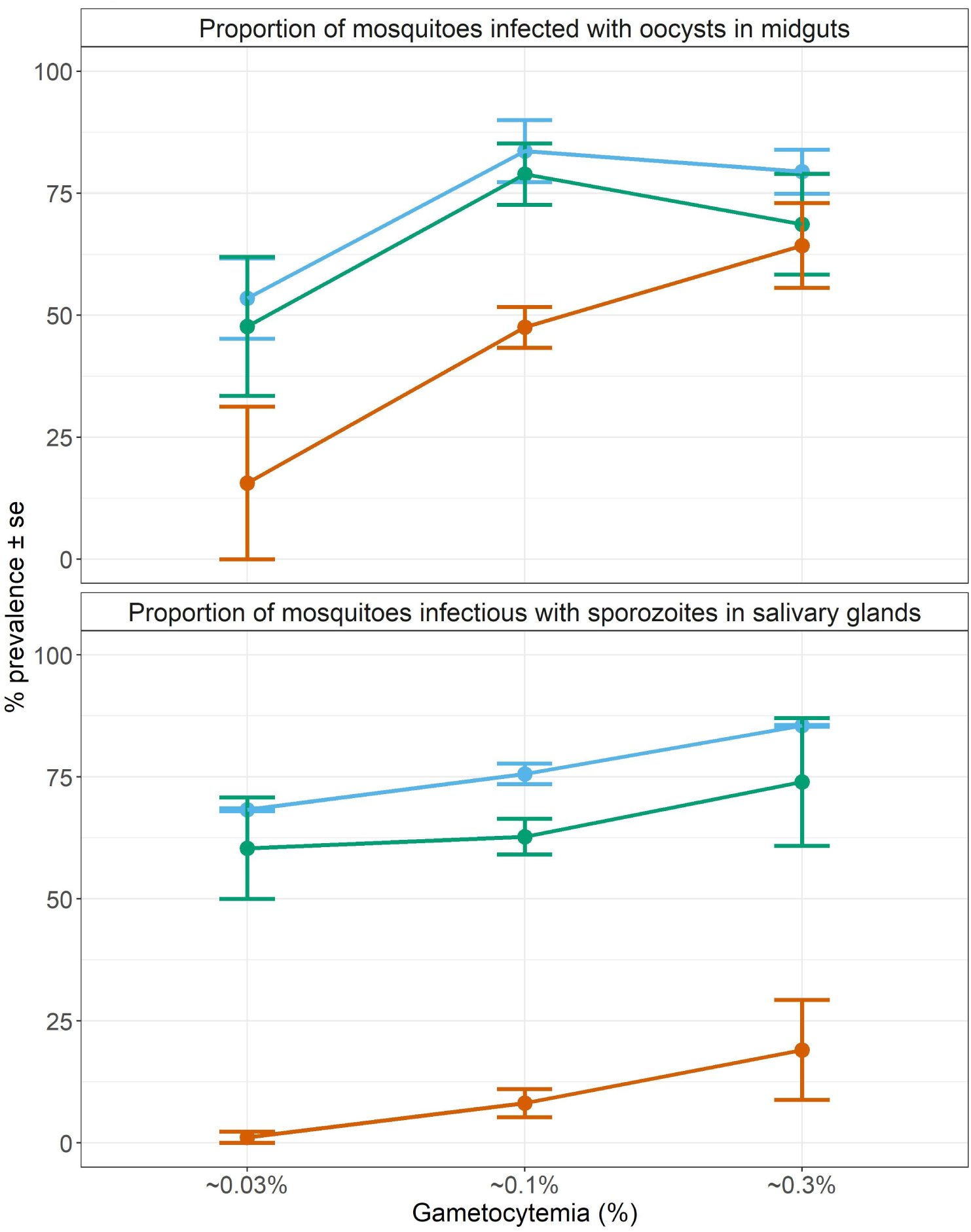
Effect of gametocyte density and temperature on oocyst prevalence in midguts (top pane) and sporozoite prevalence in the salivary glands (bottom pane). Values represents means ± standard error (se) from three biological replicates (1469 mosquitoes).

### The effects of temperature and gametocyte density on oocyst intensity

In general, the mean number of oocysts per infected mosquito midgut showed significant declines as temperatures warmed, with mosquitoes housed at 20 DTR 9°C experiencing on average the highest parasite burdens (Figure 3, Table 2). Further, increases in gametocytemia resulted in positive and linear increases in the mean number of oocysts per midgut, albeit with weak evidence for a quadratic relationship where oocyst intensity was higher at the intermediate gametocyte densities (Figure 3, Table 2 and supplementary table 3). Of the three temperatures however, only at the warmest temperature (28 DTR 9°C) was the relationship between gametocyte density and oocyst intensity clearly linear with higher densities resulting in higher burdens in mosquito midguts relative to the other two temperatures (Figure 3, Table 2). Overall, the model was able to predict a total of 57.9% of the variation, with the predictors contributing 52.2% (Table 2). Pairwise comparisons of oocyst intensity suggest clear differences in the mean number of oocysts per midgut across all three gametocyte densities at the two cooler temperatures of 20 and 24 DTR 9°C. At 28 DTR 9°C, only the highest and lowest densities differed significantly in their contribution to burdens, however, this interpretation should be taken with caution since mosquitoes in two of the three experimental replicates at this temperature showed no evidence of infection at the lowest density, which may in turn have affected the pairwise comparisons.

**Table 2:** Statistical models for oocyst intensity (infected midguts).

| Table 2 |  |  |  |  |
| --- | --- | --- | --- | --- |
|  | *Oocyst intensity* |  |  |  |
| *Predictors (reference group = 20 DTR 9°C)* | *Log-Mean* | *std. Error* | *z-value* | *p* |
| (Intercept) | 2.112 | 0.143 | 14.772 | <**0.001** |
| 24 DTR 9°C | -0.580 | 0.168 | -3.449 | **0.001** |
| 28 DTR 9°C | -1.386 | 0.185 | -7.481 | <**0.001** |
| Gametocyte density (linear trend) | 12.513 | 1.078 | 11.612 | <**0.001** |
| Gametocyte density (quadratic / “hump-shaped” trend) | -3.234 | 1.072 | -3.016 | **0.003** |
| 24 DTR 9°C / Gametocyte density (linear trend) | -2.271 | 1.634 | -1.390 | 0.165 |
| 28 DTR 9°C / Gametocyte density (linear trend) | -6.434 | 2.342 | -2.747 | **0.006** |
| 24 DTR 9°C / Gametocyte density (quadratic / “hump-shaped” trend) | -2.749 | 1.624 | -1.692 | 0.091 |
| 28 DTR 9°C / Gametocyte density (quadratic / “hump-shaped” trend) | -0.565 | 2.558 | -0.221 | 0.825 |
| **Random Effects** |  |  |  |  |
| Random variation in intercepts between temperatures within each replicate | 0.03 |  |  |  |
| Random variation in intercepts between replicates | 0.1 |  |  |  |
| Observations | 504 |  |  |  |
| Marginal R<sup>2</sup> / Conditional R<sup>2</sup> | 0.522 / 0.579 |  |  |  |
| GLMM family (“link”) | Negative binomial with variance increasing quadratically with the mean<br>(glmmTMB family= nbinom2 (link = “log”)) |  |  |  |

### The effects of temperature and gametocytemia on the distribution of parasites across mosquitoes

Simple phenomenological analyses of the oocyst distribution across individual mosquito midguts suggests strong gametocyte density- and temperature-dependent patterns (Figure 5 and supplementary figure 1). To describe how parasites are distributed across mosquitoes we used the variance to mean ratio, which is a common metric often used to estimate the amount of parasite aggregation across a population of hosts (Wilson et al., 2002). Exclusion of un-infected midguts (oocyst counts ≥ 1 in Figure 5, defined here as oocyst intensity) suggests gametocyte density alters the degree of parasite aggregation, with higher gametocyte density generally resulting in a higher degree of parasite aggregation (i.e. parasites are dispersed unevenly across infected mosquitoes, with a few mosquitoes harboring high midgut burdens and the majority of infected mosquitoes with low midgut burdens). However, temperature did modify the effect of gametocyte density on parasite distributions. At the lowest temperature (20 DTR 9°C), increases in gametocyte density led to increased variance to mean ratios that reflect fairly low parasite aggregation to higher amounts of parasite aggregation. At the intermediate temperature, parasite distributions went from randomly distributed (variance: mean ∼ 1) to higher parasite aggregation when gametocyte density increased from the lowest density treatment (0.03%). Interestingly, at the highest temperature (28 DTR 9°C), increases in gametocyte density does not affect parasite distributions across infected mosquitoes, with parasites distributed more randomly across infected mosquitoes regardless of gametocyte density (Figure 5 and supplementary figure 1).

**Figure 4:**
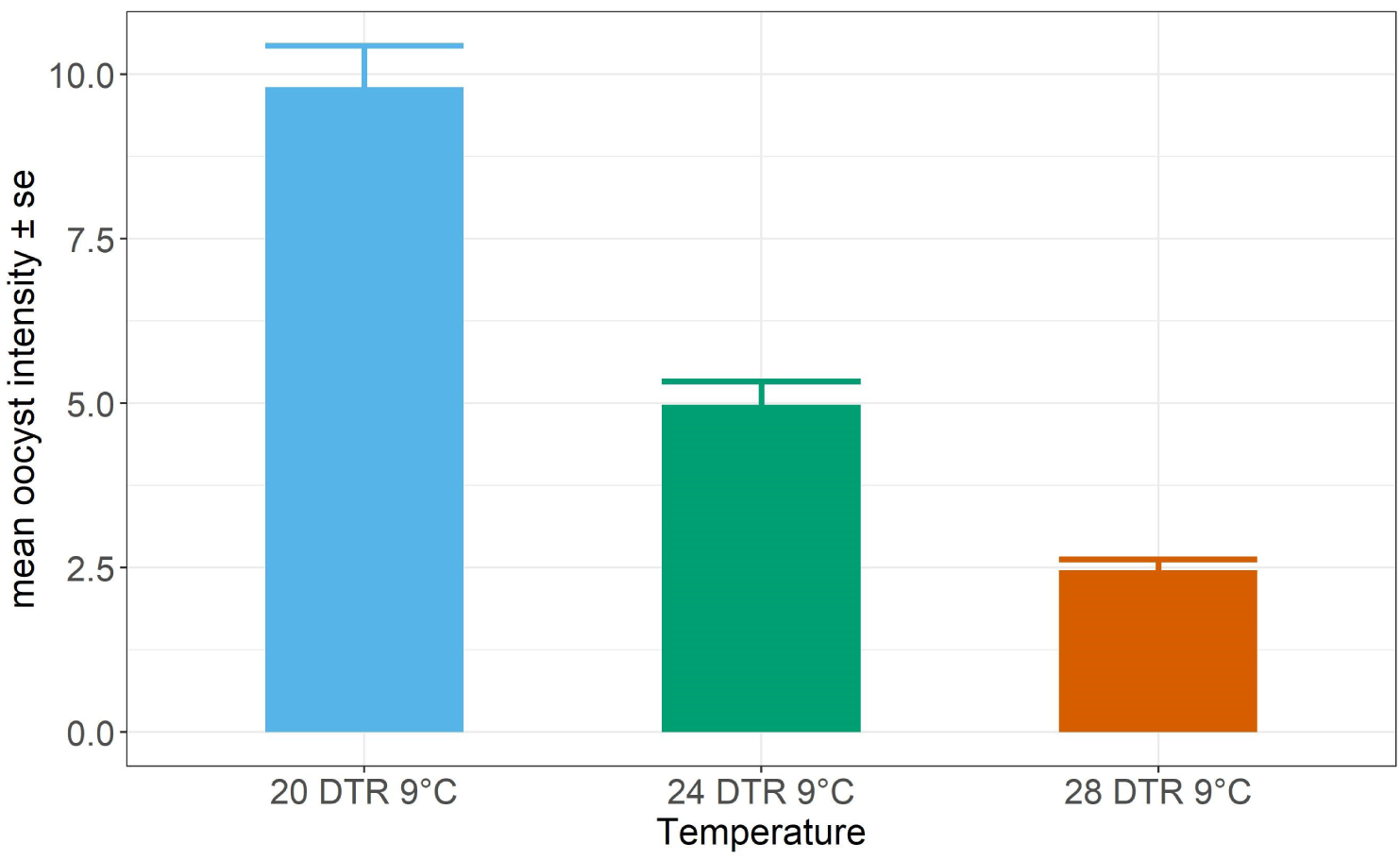
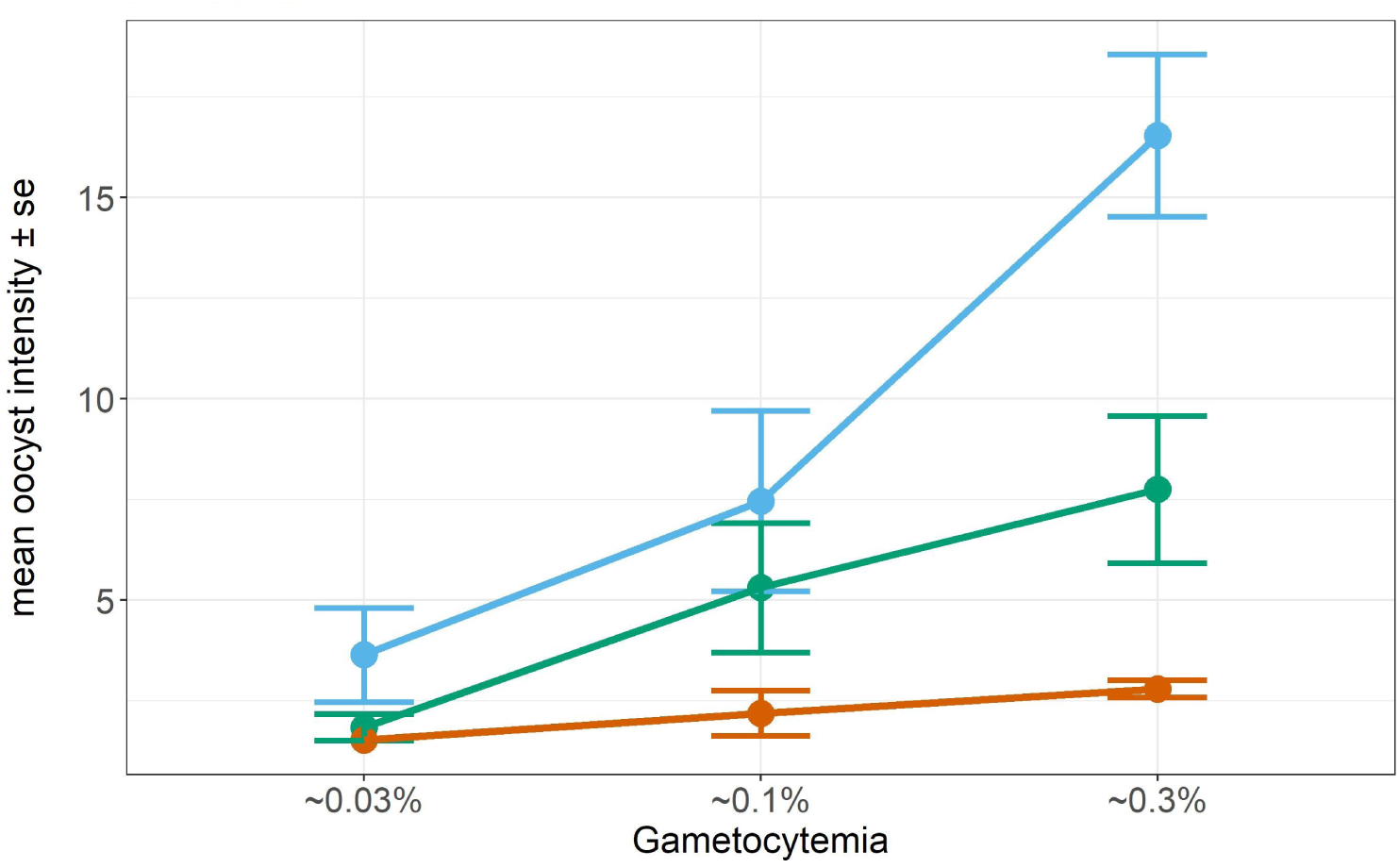
Oocyst intensities (infected midguts) across temperatures (a.) and gametocyte density (b.). Values represents means ± standard error (se) from three biological replicates (504 mosquitoes).

**Figure 5:**
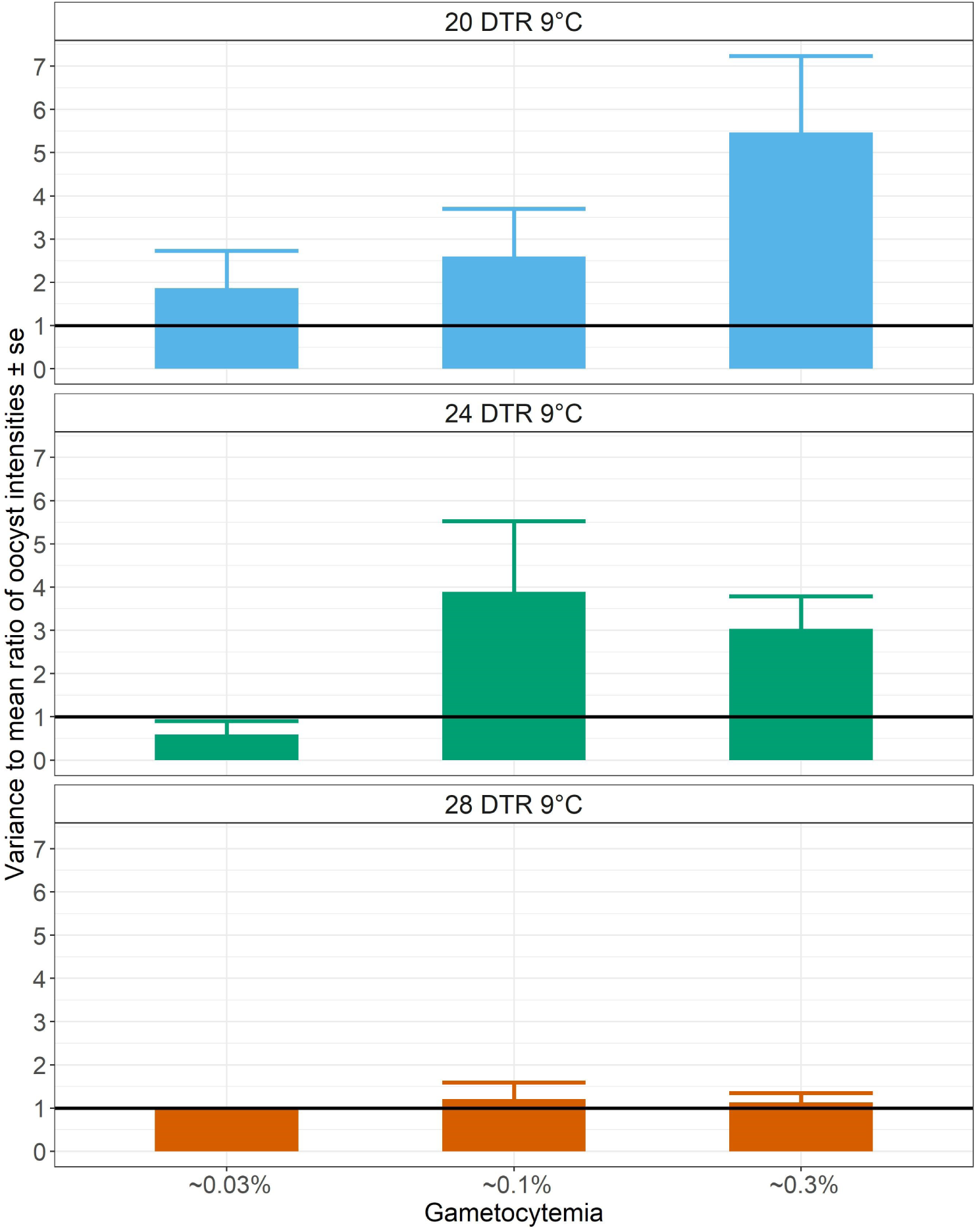
Effect of gametocytemia on the distribution of oocyst burdens across infected individual midguts at 20 DTR 9°C, 24 DTR 9°C, and 28 DTR 9°C using the variance to mean ratio (*s*^*2*^: *m*). Values represents means ± standard error (se) from three biological replicates (504 mosquitoes).

## Discussion

Overall, increasing temperatures exert a negative effect on *P. falciparum* fitness with declines in the mean proportion of mosquitoes that become infected and infectious, as well as the intensity of infection. The main effects of temperature observed in this study on vector competence fall in line with predicted temperature-vector competence relationships for malaria outlined in previous studies (Mordecai et al., 2013;Johnson et al., 2015;Shapiro et al., 2017). Further, while a relatively similar proportion of mosquitoes were infected and infectious at the cool and intermediate temperature, a significant drop was noted in the proportion of mosquitoes that become infectious at the warmest temperature. This suggests that the overall efficiency of malaria infection declines at the warmest temperature either due to a direct effect of temperature on the ability of the parasite to infect and replicate, on mosquito immune-physiology, or likely some combination of both.

The relationship between the density of transmission stages a mosquito is exposed to and metrics of vector competence, parasite burden, and parasite distribution were shaped by variation in ambient temperature. While increases in gametocyte density in general increased the proportion of mosquitoes that become infected (oocyst prevalence), the qualitative shape of this relationship varied across mosquitoes housed at different temperatures. For example, intermediate gametocytemia maximized oocyst prevalence at the cool and intermediate temperatures while high gametocytemia maximized oocyst prevalence at the warm temperature. Interestingly, the differences observed in oocyst prevalence across our temperature treatments were greatly minimized at the highest gametocytemia. At the cool and intermediate temperatures, increasing gametocyte density had a positive influence on the probability of oocysts establishing in the midgut but not on the proportion of mosquitoes that go on to become infectious, however, at the warmest temperature, a positive linear relationship was noted for the proportion infected as well as infectious. In contrast to the effect of gametocyte density on proportion of oocyst-infected mosquitoes, oocyst burdens in infected midguts showed a generally linear, positive relationship with density at the cool and intermediate temperatures. This is despite the fact that we see increases in the numbers of oocysts establishing on midguts with increasing gametocytemia, especially in mosquitoes housed at the cool and intermediate temperatures. Additionally, temperature introduced significant heterogeneity in the distribution of burdens across infected midguts in a gametocyte density-dependent manner. At the warmest temperature, parasites were randomly distributed across infected hosts regardless of initial gametocyte density, suggesting mosquitoes are more resistant to infection at this temperature in general. At the lower temperatures however, the distributions showed aggregation in a density-dependent manner suggesting that temperature and density together may expose significant variability in responses between individual mosquitoes.

Whether these effects of temperature on parasite metrics and distributions are due to direct effects of temperature on the parasite or indirect effects mediated through changes in mosquito anti-malarial responses warrants further investigation, however, our results may have several important implications for understanding malaria transmission (and control). *First*, if ambient temperature modifies the relationship between gametocyte density and transmission, it could provide an additional explanation for why the distribution patterns of allelic variants at and around the *gdv1* (gametocyte development 1) and/or *AP2-g* (apicomplexan apetela-2) locus, which are distinguished primarily by their investment in gametocytogenesis, vary seasonally and geographically (Gadalla et al., 2016;Duffy et al., 2018;Rono et al., 2018;Usui et al., 2019). *Second*, while higher temperatures could select for transmission of genotypes capable of higher gametocytogenesis, at lower temperatures the same genotype could be outcompeted due to density-dependent effects. *Third*, identifying human infectious reservoirs and evaluating their contribution to overall transmission to better target intervention efforts will also likely depend on environmental context. For instance, low-density individuals might contribute as much as children carrying high gametocyte densities in regions of the world or times of season when temperatures are cooler and thus, intervention strategies may need to cover a relatively larger proportion of the human population. This could be further compounded if these hosts also experience higher mosquito feeding rates than the younger age-groups (Goncalves et al., 2017;Guelbeogo et al., 2018). In contrast, age cohorts with higher gametocytemia might contribute more to overall transmission in warmer regions of the world or times of season.

The interactions between temperature and gametocyte density described here may have several ramifications for current transmission reducing / blocking interventions. *First*, transmission reducing interventions, such as the anti-gametocidal drug primaquine, could be much more effective in reducing transmission of artemisinin resistant parasites in warmer environments or times of season than in cooler seasons or geographic regions. *Second*, thermal variation in the field could have implications for current vaccine design and testing pipelines where the rate of reduction in oocyst “intensity” (the terminological equivalent of abundance in this study) is the primary method of evaluation (Bompard et al., 2017). For instance, more effective antibody responses may be required at lower temperatures where oocyst burdens and sporozoite prevalence are highest. *Third*, if transmission blocking vaccines are imperfect as early evidence suggests (Sagara et al., 2018), intermediate gametocytemia in some environmental contexts might actually boost transmission. This suggests that at lower temperatures, density-dependent constraints such as competition between parasites for limited resources or temperature effects on mosquito anti-malarial responses may facilitate “transmission enhancement” in mosquitoes exposed to the reduced gametocyte densities. Such transmission enhancement is not without precedent and has also been observed in presence of low levels of anti-gametocytic antibodies (Stone et al., 2018).

There are some limitations to the current study. *First*, the statistical analyses clearly indicate heterogeneity in the data, especially for prevalence for which the models accounted for <50% of the variation, but also oocyst burdens to a certain extent. Indeed, while it is not possible to rule out technical variations in the serial dilutions across each experimental replicate, logistical constraints associated with the inherently low-throughput nature of SMFAs meant only three gametocyte densities could be assessed at each temperature, which may have obscured clearer associations. *Second*, while it is possible that the gametocyte densities tested in the current study are more in line with laboratory rather than field-based studies (Koepfli and Yan, 2018), our key conclusion should still hold – at cooler temperatures, successful transmission should still be achieved at even lower densities. For instance, at the two cooler temperatures, mean prevalence in either organ was ≥50%. *Third*, vector competence was measured with a “non-native” vector species-parasite genotype with the mosquito originating from South Asia and the parasite thought to be of African descent (Molina-Cruz et al., 2015). However, in addition to good overall correlation in vector competence of *An. stephensi* and the African vector *An. gambiae* (Murdock et al., 2016;Eldering et al., 2017), the study could prove to be timely considering the recent migration of *An. stephensi* to Eastern Africa (Takken and Lindsay, 2019). *Fourth*, while only one regime was tested here (DTR 9°C), it is possible that the magnitude of fluctuations may result in different parasite fitness phenotypes and collectively argues for investigating the underlying relationships in a fluctuating rather than constant temperature environment (Paaijmans et al., 2010;Blanford et al., 2013;Murdock et al., 2016). *Fifth*, we did not measure sporozoite burdens in the salivary glands. Intensity of *P. falciparum* oocysts in the midguts have been shown to positively correlate with sporozoite numbers in the salivary glands (Stone et al., 2013;Miura et al., 2019). However, previous work in rodent malaria systems have demonstrated negative relationships between oocyst burden and the number of sporozoites per oocyst, as well as the rate at which sporozoites invade the salivary glands (Pollitt et al., 2013;Moller-Jacobs et al., 2014). Our results argue for the consideration of field relevant environmental contexts as an important determinant of density-dependent effects on sporozoite prevalence and burdens.

In this study, we demonstrate that field relevant variation in ambient temperature alters the relationship between density of transmission stages and infection outcomes in the *An. stephensi* – *P. falciparum* system. In particular, we observed qualitatively different effects of temperature on the relationship between gametocytemia and parasite prevalence (oocyst and sporozoite), burdens (oocyst intensity), and distribution across exposed mosquitoes. We emphasize that the role of variation in field relevant factors like temperature in shaping the relationship between gametocytemia and mosquito infection is currently under appreciated, with potentially important ramifications for understanding natural transmission and malaria control. Thus, it is imperative to begin incorporating field relevant sources of variation into our mechanistic understanding of malaria transmission. This study represents an important step in this direction.

## Acknowledgements

Funding for this work was provided by the University of Georgia and the NIH (5R01AI110793-04). The authors would also like to thank members of the Murdock lab for their patience and support.

## Author contributions

AKP designed the study, collected and analyzed the data and wrote the manuscript with critical input from CCM. JCS contributed to the mosquito husbandry and helped AKP collect data. MBT provided comments and suggestions during the project and on the manuscript. All authors read and approved the final version of the manuscript.

---

**Supplementary table 1:** Statistical models for prevalence of oocysts and sporozoites in midguts and salivary glands for each temperature and DTR. ^#^ Sporozoite prevalence data from one biological replicate was not available. ^+^ Sporozoite prevalence data from the lowest density for one biological replicate was not available.

|  | 20 DTR 9°C |  |  |  | 24 DTR 9°C |  |  |  | 28 DTR 9°C |  |  |  |
| --- | --- | --- | --- | --- | --- | --- | --- | --- | --- | --- | --- | --- |
| *Predictors* | *Log-Odds* | *std. Error* | *Z value* | *p* | *Log-Odds* | *std. Error* | *Z value* | *p* | *Log-Odds* | *std. Error* | *Z value* | *p* |
| (Intercept) | 1.021 | 0.146 | 6.995 | **<0.001** | 0.729 | 0.274 | 2.658 | **0.007** | -0.569 | 0.425 | -1.338 | 0.181 |
| Sporozoite prevalence | 0.199 | 0.232 | 0.858 | 0.390 | -0.02 | 0.282 | -0.073 | 0.942 | -2.04 | 0.575 | -3.547 | **<0.001** |
| Linear trend over parasite density | 1.07 | 0.551 | 1.942 | 0.052 | 0.558 | 0.75 | 0.743 | 0.457 | 4.241 | 1.287 | 3.293 | **<0.001** |
| Quadratic trend over parasite density | -1.947 | 0.579 | -3.363 | **<0.001** | -1.996 | 0.792 | -2.519 | **0.012** | -2.707 | 1.316 | -2.056 | **0.039** |
| Sporozoite prevalence * linear trend over parasite density | 0.53 | 0.926 | 0.573 | 0.566 | 0.483 | 1.161 | 0.416 | 0.677 | -1.015 | 2.268 | -0.447 | 0.654 |
| Sporozoite prevalence * quadratic trend over parasite density | 1.771 | 0.875 | 2.024 | **0.042** | 2.253 | 1.164 | 1.935 | 0.052 | 0.501 | 2.258 | 0.222 | 0.824 |
| **Random Effects** |  |  |  |  |  |  |  |  |  |  |  |  |
| Random variation in intercepts between the three biological replicates | 0.00 |  |  |  | 0.15 |  |  |  | 0.26 |  |  |  |
| Number of mosquitoes sampled | 449<sup>#</sup> |  |  |  | 483<sup>+</sup> |  |  |  | 537 |  |  |  |

**Supplementary table 2:** Pairwise comparisons of means predicted by the model in Table 1 in the main text for oocyst and sporozoite prevalence.

| Oocyst prevalence (midguts) |  |  |  |  |
| --- | --- | --- | --- | --- |
| Temperature | Pairwise comparisons<br>Gametocytemia 1 vs gametocytemia 2 | estimate (log odds) | std. Error | *p*-value |
| 20 DTR 9°C | ~0.03% vs. ~0.1% | -0.286 | 0.092 | **0.012** |
| 20 DTR 9°C | ~0.03% vs. ~0.3% | -0.233 | 0.095 | 0.053 |
| 20 DTR 9°C | ~0.1% vs. ~0.3% | 0.053 | 0.080 | 0.786 |
| 24 DTR 9°C | ~0.03% vs. ~0.1% | -0.277 | 0.093 | **0.015** |
| 24 DTR 9°C | ~0.03% vs. ~0.3% | -0.172 | 0.098 | 0.200 |
| 24 DTR 9°C | ~0.1% vs. ~0.3% | 0.105 | 0.090 | 0.477 |
| 28 DTR 9°C | ~0.03% vs. ~0.1% | -0.345 | 0.096 | **0.003** |
| 28 DTR 9°C | ~0.03% vs. ~0.3% | -0.507 | 0.093 | **<.0001** |
| 28 DTR 9°C | ~0.1% vs. ~0.3% | -0.162 | 0.100 | 0.249 |
| Sporozoite prevalence (salivary glands) |  |  |  |  |
| Temperature | Pairwise comparisons<br>Gametocytemia 1 vs gametocytemia 2 | estimate (log odds) | std. Error | *p*-value |
| 20 DTR 9°C | ~0.03% vs. ~0.1% | -0.069 | 0.111 | 0.812 |
| 20 DTR 9°C | ~0.03% vs. ~0.3% | -0.165 | 0.104 | 0.266 |
| 20 DTR 9°C | ~0.1% vs. ~0.3% | -0.097 | 0.097 | 0.585 |
| 24 DTR 9°C | ~0.03% vs. ~0.1% | -0.005 | 0.120 | 0.999 |
| 24 DTR 9°C | ~0.03% vs. ~0.3% | -0.120 | 0.116 | 0.560 |
| 24 DTR 9°C | ~0.1% vs. ~0.3% | -0.116 | 0.102 | 0.503 |
| 28 DTR 9°C | ~0.03% vs. ~0.1% | -0.069 | 0.043 | 0.258 |
| 28 DTR 9°C | ~0.03% vs. ~0.3% | -0.134 | 0.059 | 0.074 |
| 28 DTR 9°C | ~0.1% vs. ~0.3% | -0.066 | 0.066 | 0.582 |

**Supplementary table 3:**
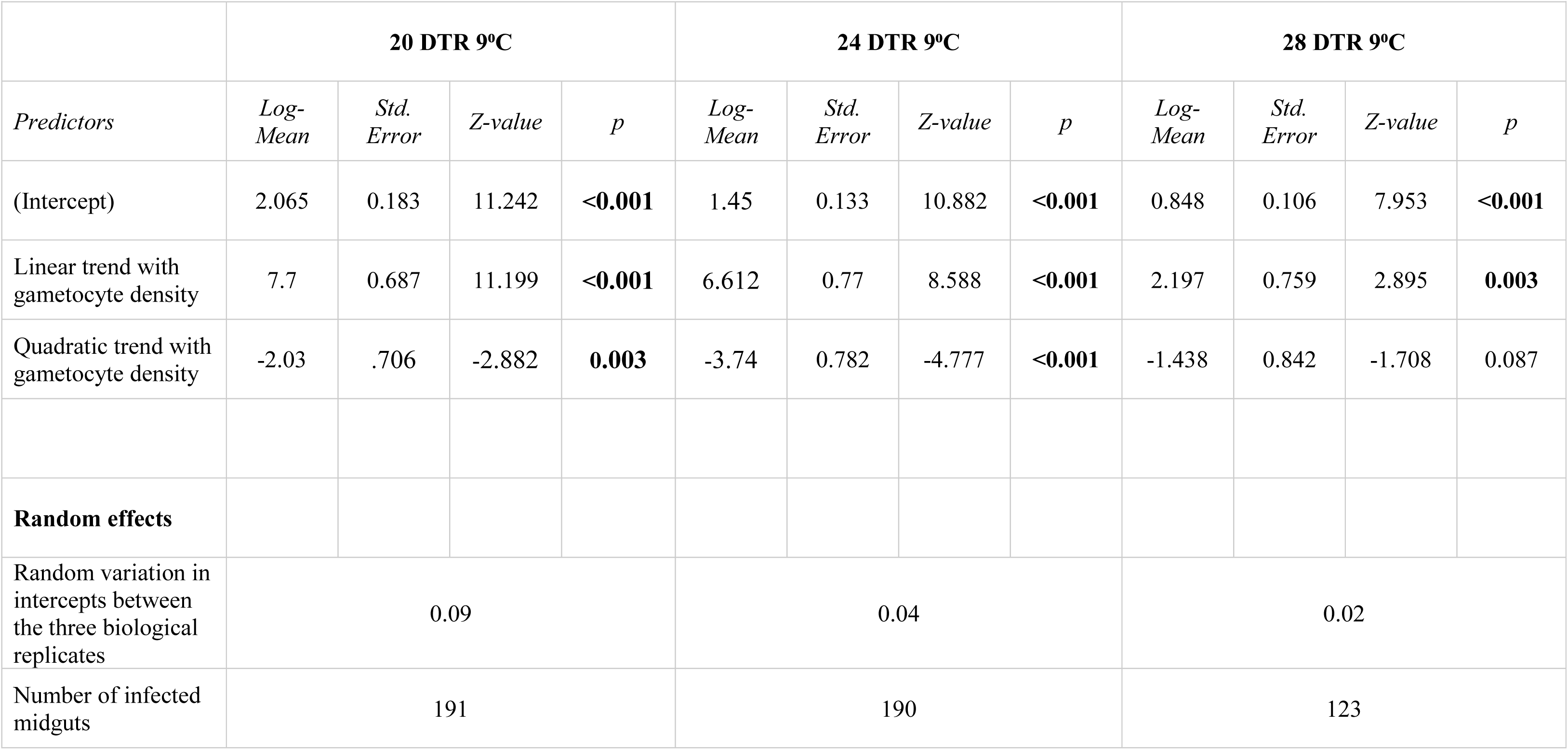
Statistical models for oocyst intensity (infected midguts) for each temperature.

|  | 20 DTR 9°C |  |  |  | 24 DTR 9°C |  |  |  | 28 DTR 9°C |  |  |  |
| --- | --- | --- | --- | --- | --- | --- | --- | --- | --- | --- | --- | --- |
| *Predictors* | *Log-Mean* | *Std. Error* | *Z-value* | *p* | *Log-Mean* | *Std. Error* | *Z-value* | *p* | *Log-Mean* | *Std. Error* | *Z-value* | *p* |
| (Intercept) | 2.065 | 0.183 | 11.242 | **<0.001** | 1.45 | 0.133 | 10.882 | **<0.001** | 0.848 | 0.106 | 7.953 | **<0.001** |
| Linear trend with gametocyte density | 7.7 | 0.687 | 11.199 | **<0.001** | 6.612 | 0.77 | 8.588 | **<0.001** | 2.197 | 0.759 | 2.895 | **0.003** |
| Quadratic trend with gametocyte density | -2.03 | .706 | -2.882 | **0.003** | -3.74 | 0.782 | -4.777 | **<0.001** | -1.438 | 0.842 | -1.708 | 0.087 |
| **Random effects** |  |  |  |  |  |  |  |  |  |  |  |  |
| Random variation in intercepts between the three biological replicates | 0.09 |  |  |  | 0.04 |  |  |  | 0.02 |  |  |  |
| Number of infected midguts | 191 |  |  |  | 190 |  |  |  | 123 |  |  |  |

**Supplementary table 4:** Pairwise comparisons of means predicted by the model in Table 2 in the main text for oocyst abundance and intensity.

| Oocyst intensity (infected midguts) |  |  |  |  |
| --- | --- | --- | --- | --- |
| Temperature | Pairwise comparisons<br>Gametocytemia 1 vs gametocytemia 2 | estimate (log odds) | std. Error | *p*-value |
| 20 DTR 9°C | ~0.03% vs. ~0.1% | -3.681 | 0.83 | **<.0001** |
| 20 DTR 9°C | ~0.03% vs. ~0.3% | -12.253 | 2.049 | **<.0001** |
| 20 DTR 9°C | ~0.1% vs. ~0.3% | -8.572 | 1.73 | **<.0001** |
| 24 DTR 9°C | ~0.03% vs. ~0.1% | -3.07 | 0.637 | **<.0001** |
| 24 DTR 9°C | ~0.03% vs. ~0.3% | -5.61 | 1.009 | **<.0001** |
| 24 DTR 9°C | ~0.1% vs. ~0.3% | -2.54 | 0.838 | **0.0072** |
| 28 DTR 9°C | ~0.03% vs. ~0.1% | -0.975 | 0.44 | 0.0694 |
| 28 DTR 9°C | ~0.03% vs. ~0.3% | -1.533 | 0.475 | **0.0038** |
| 28 DTR 9°C | ~0.1% vs. ~0.3% | -0.558 | 0.403 | 0.3502 |

**Supplementary figure 1:**
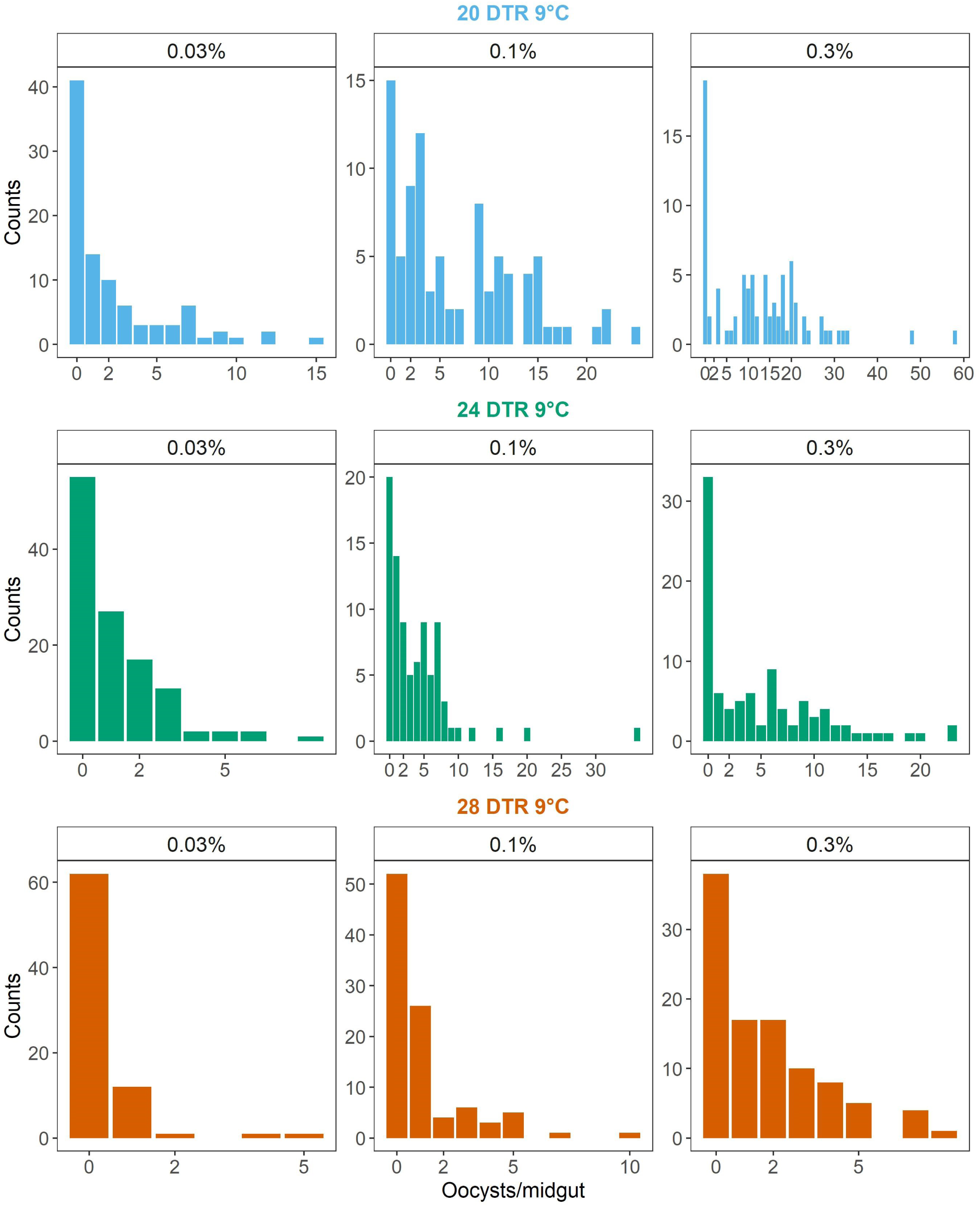
Effect of gametocytemia on the distribution of oocyst burdens across all midguts (infected and un-infected) at 20 DTR 9°C (blue, top panes), 24 DTR 9°C (green, middle panes) and 28 DTR 9°C (red, bottom panes). Gametocyte densities are indicated in the strips above each plot. Values represent data from 839 mosquito midguts from three biological replicates.

